# Hypothalamic prostaglandins facilitate recovery from hypoglycemia but exacerbate recurrent hypoglycemia in mice

**DOI:** 10.1101/2024.06.24.600540

**Authors:** Takashi Abe, Shucheng Xu, Yuki Sugiura, Yuichiro Arima, Takahiro Hayasaka, Ming-Liang Lee, Taiga Ishimoto, Yudai Araki, Samson Ngurari, Ziwei Niu, Norifumi Iijima, Sabrina Diano, Chitoku Toda

## Abstract

The hypothalamus monitors blood glucose levels and regulates glucose production in the liver. In response to hypoglycemia, glucose-inhibited (GI) neurons trigger counter-regulatory responses (CRRs), which stimulate the release of glucagon, epinephrine, and cortisol to elevate blood glucose. Recurrent hypoglycemia (RH), however, reduces the effectiveness of these CRRs. This study examined the role of hypothalamic prostaglandins in glucose recovery during acute hypoglycemia and RH. Using imaging mass spectrometry and liquid chromatography/mass spectrometry, phospholipid and prostaglandin levels in the hypothalamus of C57BL mice were increased following insulin or 2-deoxy-glucose administration. Ibuprofen, a nonsteroidal anti-inflammatory drug (NSAID), was infused into the ventromedial hypothalamus (VMH) to analyze its effect on glucose production during hypoglycemia, revealing that prostaglandin inhibition decreased glucagon secretion. Additionally, RH-treated mice decreased glucagon release and glucose production during hypoglycemia. Inhibiting prostaglandin production via short-hairpin RNA against cytosolic phospholipase A2 (cPLA2) in the hypothalamus restored CRRs diminished by RH via increasing glucagon sensitivity. These findings suggest that hypothalamic prostaglandins play a critical role in glucose recovery from acute hypoglycemia by activating VMH neurons and are also crucial for the attenuation of CRRs during RH.

**Graphical Abstract:** 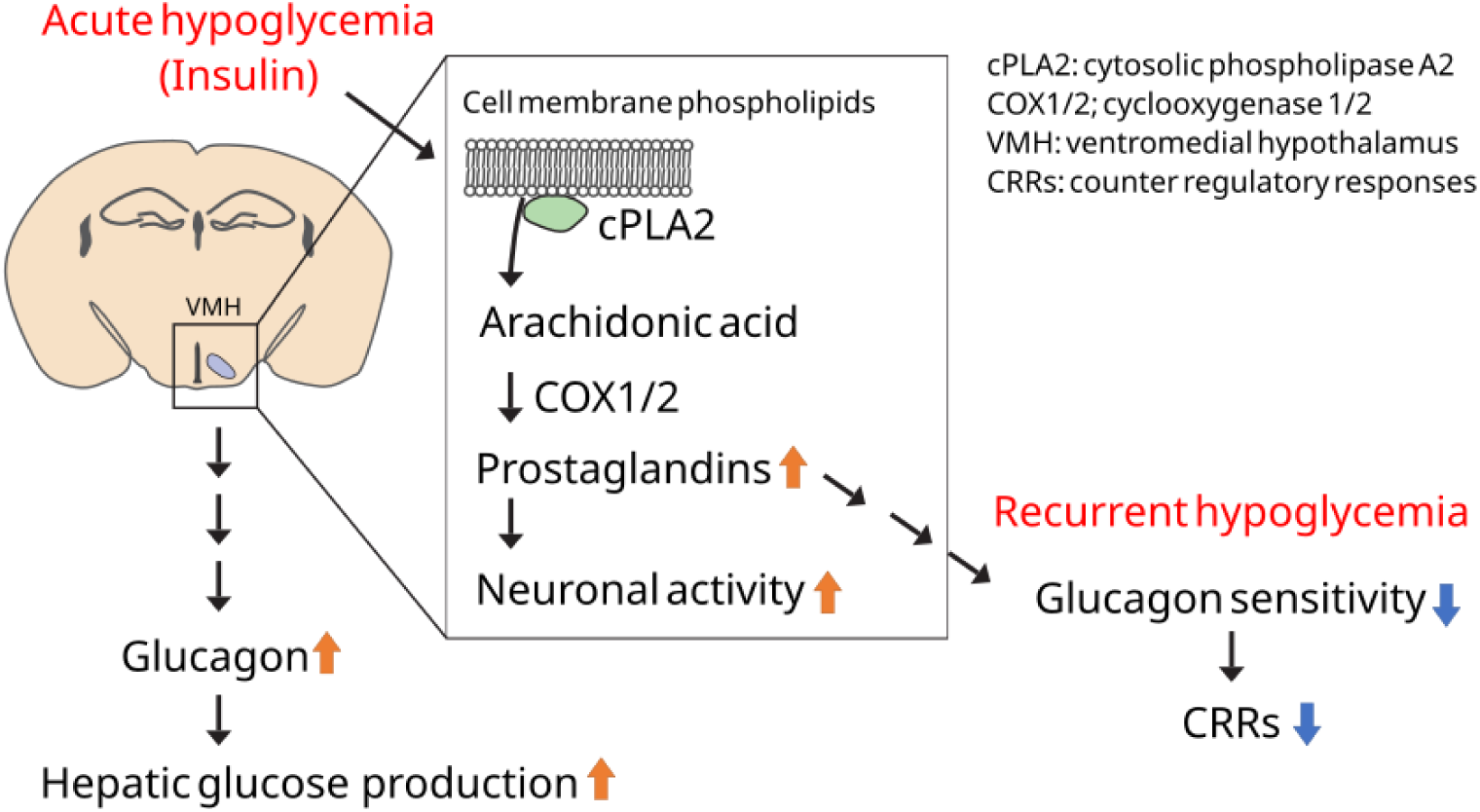

**Article Highlights:**

- Insulin decreases arachidonic acid-containing phospholipids and increases prostaglandin production in the hypothalamus.
- Prostaglandin in the hypothalamus increases glucagon secretion and glucose production during hypoglycemia.
- Recurrent hypoglycemia decreases glucagon secretion and glucose production during hypoglycemia.
- Prostaglandin in the hypothalamus during recurrent hypoglycemia decreases glucagon sensitivity and glucose production.

## Introduction

Hypoglycemia, a common side effect of diabetes medications, can lead to severe consequences, including lightheadedness, fainting, and even death. Continuous glucose monitoring systems have revealed that hypoglycemic events occur more frequently than previously thought (1). Recurrent hypoglycemia (RH) is defined as experiencing multiple episodes of hypoglycemia, which alters the body’s response to hypoglycemia and increases the risk of severe hypoglycemia (2). Thus, hypoglycemia-related deaths worldwide are assumed to be 4.49 per 1,000 total diabetes deaths (3). Additionally, RH increases the risk of dementia (4). The increasing prevalence of hypoglycemia-related deaths and dementia underscores the need for a deeper understanding of its underlying mechanisms.

To counteract hypoglycemia, the body initiates counter-regulatory responses (CRRs) involving the secretion of glucagon, corticosteroids, and catecholamines to stimulate hepatic glucose production and restore blood glucose levels (5). The brain regulates CRRs using two glucose-sensing neurons: glucose-inhibited (GI) and glucose-excited (GE) neurons. GI neurons, activated by hypoglycemia, enhance glucose production by modulating CRRs (6). Various neuronal populations, including neuropeptide Y neurons in the arcuate nucleus of the hypothalamus (ARC), hypocretin/orexin neurons in the lateral hypothalamus, and CCK- or leptin receptor-expressing neurons in the parabrachial nucleus (PBN), exhibit GI characteristics (7–10). Within the ventromedial hypothalamus (VMH), pituitary adenylate-cyclase-activating polypeptide, glucokinase, nitric oxide synthase 1 (NOS1), cholecystokinin (CCK)-B receptor, and anoctamin 4 channel-positive neurons are also GI and contribute to glucose elevation (11–14). When hypoglycemia occurs, intracellular ATP levels drop, and the AMP/ATP ratio increase activates AMP-activated protein kinase and produces nitric oxide to depolarize GI neurons (15). When glucose is unavailable in the brain, neurons utilize lactate as an alternative fuel. Thus, a lactate infusion into the VMH during hypoglycemia attenuates CRRs (16). In contrast, GE neurons regulate insulin sensitivity in peripheral tissues during hyperglycemia (17). GE neuron also affects CRRs during hypoglycemia via the K_ATP_ channel (18).

Recurrent hypoglycemia (RH) attenuates CRRs (19). Central and peripheral mechanisms have been reported (20,21). In the central nervous system, RH affects GE and GI neurons’ sensitivity to glucose (22,23). Lactate production from glycogen and fatty acid oxidation in astrocytes in the hypothalamus also involves the development of RH (24). Lactate is utilized as an alternative fuel in the brain after RH (25). However, the mechanism by which the hypothalamus adapts in response to RH has not been fully understood.

Non-steroidal anti-inflammatory drugs (NSAIDs) are known to develop hypoglycemia by increasing insulin release through the K_ATP_ channel in pancreatic beta cells (26). NSAIDs also decrease gluconeogenesis by affecting hepatocytes (27). Recently, we have reported that high blood glucose increases prostaglandin production via phospholipase A2 (PLA2) in the hypothalamus and activates GE neurons (17). Similar to our previous report, insulin-induced hypoglycemia inhibits arachidonic acid metabolism of phospholipids in the brain (28). Hypoxia and hypoglycemia activate PLA2, increase the release of arachidonic acid, and increase cFos in neuroblastoma cells (29). From these reports, we hypothesized that hypoglycemia increases the production of prostaglandins through PLA2-mediated arachidonic acid release, which alters the activities of GI and GE neurons. However, the role of hypothalamic prostaglandin in CRRs during hypoglycemia is unclear.

This study aimed to investigate hypothalamic prostaglandin’s role in CRRs during acute hypoglycemia and RH. Our findings suggest that hypoglycemia increases prostaglandin production in the hypothalamus, which is essential for activating VMH neurons and stimulating glucagon secretion. Furthermore, inhibiting phospholipid metabolism through cPLA2 shRNA in the hypothalamus improves the attenuation of CRRs during RH.

## Research Design and Methods

### Animals

C57L/6N male and female mice (The Jackson Laboratory Japan, Yokohama, Japan) were kept at room temperature with a 12h/12h light and dark time cycle. The mice had free access to water and a regular chow diet (Nosan Corporation, Yokohama, Japan). Experiments were carried out in the Experimental Animal Facility of Hokkaido University (AAALAC Accredited) and Kumamoto University. Mice cages were changed once a week, and mice care was performed according to the Animal Care and Use Committee guidelines of Hokkaido University and Kumamoto University.

### Recurrent hypoglycemia

C57L/6N male and female mice received i.p. injection of insulin (2.5 U/kg, Humarin R, Eli Lilly Japan, Hyogo, Japan) at 1 pm. Blood glucose levels were measured by a handy glucose meter (Nipro FreeStyle, Nipro, Osaka, Japan). When blood glucose reached 40 mg/dL, glucose (3 g/kg) was injected i.p. or subcutaneously to rescue from hypoglycemia. The same procedure was repeated 5 times.

### Imaging mass spectrometry

C57BL male mice received i.p. injection of insulin (1 U/kg, 30 min), 2DG (2 g/kg, 60 min, Sigma-Aldrich, St. Louis, MO), or saline, after which they were sacrificed using CO_2_ asphyxiation. The mice brains were immediately placed into ice-cold saline, embedded in 2% sodium carboxymethyl cellulose solution, and frozen with liquid nitrogen. The 10-μm brain sections were prepared by cryostat and immediately mounted onto an indium-tin-oxide-coated glass slide (Bruker Daltonics, Bremen, Germany). The sections on the glass slides were immediately dried and stored at −20°C until imaging mass spectrometry analysis.

The brain sections were sprayed with 9-aminoacridine matrix (10 mg/mL in 70% ethanol, Sigma-Aldrich) and installed into a matrix-assisted laser desorption/ionization (MALDI)-time-of-flight (TOF)/TOF system using ultrafleXtreme (Bruker Daltonics). The brain sections were irradiated by a smart beam (Nd: YAG laser, 355-nm wavelength), with a 25-μm irradiation pitch. The laser had a repetition frequency of 2000 Hz and mass spectra were obtained in the range of m/z 200–1200 in negative-ion mode. The m/z values from previous reports were used to label each lipid, phospholipid, and lyso species (17). All ion images were reconstructed with total ion current (TIC) normalization by flexImaging (Bruker Daltonics) and transferred to ImageJ software after modifying the grayscale. The areas of the VMH were identified by 4’,6-diamidino-2-phenylindole (DAPI) staining in the other brain sections, and the brightness of mass spectrometry signals in the VMH and ARC were calculated as intensity. Phosphatidyl-choline and phosphatidyl-glycerol were measured in the adjacent tissue sections with positive-ion mode. However, the mass spectra after MS/MS were inconclusive enough to allow a reliable data assignment, especially for fatty acid components.

### Quantification of PGs

C57BL male mice received i.p. injection of insulin (1 U/kg, 30 min), 2DG (2 g/kg, 60 min), or saline, after which they were sacrificed using CO_2_ asphyxiation. The hypothalamus was collected and immediately frozen in liquid nitrogen. The tissue was homogenized with 500 μL of methanol: formic acid (100:0.2) containing an internal standard consisting of a mixture of deuterium-labeled PGs using microtip sonication. The samples were subjected to solid phase extraction using an Oasis HLB cartridge (5 mg; Waters, Milford, MA). Samples were diluted with water: formic acid (100:0.03) to give a final methanol concentration of ∼20% by volume, applied to preconditioned cartridges, and serially washed with water: formic acid (100:0.03), water: ethanol: formic acid (90:10:0.03), and petroleum ether. Samples were eluted with 200 μL of methanol: formic acid (100:0.2). The filtrate was concentrated with a vacuum concentrator (SpeedVac, Thermo Fisher Scientific, Waltham, MA). The concentrated filtrate was dissolved in 50 μL of methanol and used for liquid chromatography/mass spectrometry (LC-MS).

The PGs in the hypothalamus were quantified by liquid chromatography/mass spectrometry. A triple-quadrupole mass spectrometer equipped with an electrospray ionization (ESI) ion source (LCMS-8040, Shimadzu Corporation, Kyoto, Japan) was used in the positive and negative-ESI and selective reaction monitoring modes. The samples were resolved on a reversed-phase column (Kinetex C8, 2.1 × 150 mm, 2.6 μm, Phenomenex, Torrance, CA) using a step gradient with mobile phase A (0.1% formic acid) and mobile phase B (acetonitrile). The gradient of mobile phase B concentration was programmed as 10% (0 min)−25% (5 min)−35% (10 min)−75% (20 min)−95% (20.1 min)−95% (28 min)−10% (28.1 min)−10% (30 min), at a flow rate of 0.4 mL/min and a column temperature of 40°C.

### Cannula implantation and adeno-associated virus (AAV) injection

C57BL male mice were anesthetized with ketamine (100 mg/kg) and xylazine (10 mg/kg) or a combination anesthetic (0.3 mg/kg of medetomidine, 4.0 mg/kg of midazolam, and 5.0 mg/kg of butorphanol). The mice were placed on the stereotaxic instrument (Narishige, Tokyo, Japan). The mice skulls were opened with a Cranial drill (dental drill). A stainless steel cannula (Plastics One, P1 Technologies, VA, USA) was inserted into the VMH using the following coordinates: an anterior-posterior (AP) direction; −1.5 (1.5 mm posterior to the bregma), lateral (L): ±0.4 (0.4 mm lateral to the bregma), dorsal-ventral (DV): −5.7 (5.7 mm below the bregma on the surface of the skull). The interval between the two cannulae was 0.8 mm, so there would be 0.4 mm on both sides. For intracerebroventricular (i.c.v.) injection, AP: − 0.3, L: 1.0, DV: −2.0. Cannulae were secured on the skulls with cyanoacrylic glue and the exposed skulls were covered with dental cement. For AAV injection, the mice were injected in both sides of the VMH with a maximum of 0.3 µL AAV9-RFP-U6-m-PLA2G4A-shRNA (1.0×10^12^ GC/mL, shAAV-268768, Vector Biolabs, PA, USA) or AAV9-U6-shRNA (scrumble, SCRM)-EF1a-GFP (1.0×10^12^ GC/mL, SL100894, Signagen Laboratories, MD, USA) using the following coordinates: AP: − 1.5, L: ± 0.5, DV: − 5.7. Open wounds were sutured after the viral injection. The mice were allowed to recover for 5–7 days before experiments were started.

Quantitative real-time-PCR was performed to assess the efficiency of shPLA2 in the hypothalamus. Mice were anesthetized with isoflurane and euthanized by cervical dislocation, and their brains were collected immediately, after which the hypothalamus was trimmed using a brain matrix. The hypothalamus was homogenized with Template Prepper (Nippon Gene, Tokyo, Japan), and the extracted RNA was reverse-transcribed using Oligo(dT)18 primer (Thermo Fisher Scientific) and M-MLV Reverse Transcriptase (Promega, Madison, WI). *Pla2g4a* and *Actin B* cDNA levels in the hypothalamus were measured by ΔΔCt method using SYBR Green reagent (Bio-Rad, Hercules, CA). The information on primers is shown in table S1. Quantitative PCR was performed with diluted cDNAs in a 10 µl reaction volume in duplicate.

### Assessment of cytosolic- or secretory-phospholipase A2 activity

C57BL male mice fed *ad libitum* were i.p. injected with either insulin (1 U/kg) or saline. The mice were sacrificed 30 min after the injection using CO_2_ asphyxiation and a -1.0 to -2.0 mm coronal section from the bregma was excised using the matrix, and the VMH and ARC areas were collected and stored at −80°C until use. Tissues were homogenized and centrifuged at 10,000 × *g* for 15 min at 4°C and supernatants were collected. The activity of cytosolic- or secretory-phospholipase A2 was measured according to the procedures described in the kit manuals (Abcam, Cambridge, UK). Phospholipases A1 (Invitrogen, Waltham, MA), phospholipase C (Sigma-Aldrich, St. Louis, MO), and phospholipase D (Sigma) were also measured according to the manufacture’s instruction. Phospholipase activity was normalized to protein concentration in each sample which is measured using bicinchoninic acid assay kit (Nacalai Tesque, Kyoto, Japan).

### 2DG-induced hyperglycemia and food intake

Ibuprofen (50 μg in 0.5 μL) or vehicle [20% dimethyl sulfoxide (DMSO) in phosphate-buffered saline (PBS)] was injected through the i.c.v cannulae in C57BL/6N mice in fed state 30 min before the i.p. injection of 2-DG (300 mg/kg). Blood glucose levels were measured by the handheld glucometer (Nipro FreeStyle, Nipro, Osaka, Japan) at -30, 0, 15, 30, 60, 90, and 120 min. For food intake, i.c.v. injection of ibuprofen and i.p. injection of 2-DG were performed as described above. The reduced weight of food was measured at time points of 0, 15, 30, 60, 90, 120, 150, and 180 min.

### Hyperinsulinemic-hypoglycemic clamp

C57BL male mice were anesthetized with pre-mixed ketamine (100 mg/kg) and xylazine (10 mg/kg) or a combination anesthetic (0.3 mg/kg of medetomidine, 4.0 mg/kg of midazolam, and 5.0 mg/kg of butorphanol). Polyethylene catheters were implanted into the right carotid artery and jugular vein. The tubes were inserted subcutaneously and protruded from the neck skin (30). The mice were allowed to recover for 3–5 days, and tubes were flushed with heparinized saline daily.

The mice were fasted for 4 h and experiments were initiated in a free-moving condition. Ibuprofen (250 μM, 0.2 μL) or vehicle (5% DMSO in PBS) was injected bilaterally into the VMH 15 min before measuring initial blood glucose levels. After measuring initial blood glucose levels (t = −15 and 0 min), a bolus of insulin (2.5 mU/kg) was injected through the jugular vein and insulin was continuously infused (1 mU/kg/min). Blood glucose levels were measured from arterial blood or tail vein every 5–10 min. Infusion of 30% glucose was performed at a variable rate via the jugular vein catheter to maintain a blood glucose level at 50 mg/dL. Erythrocytes in withdrawn blood were suspended in sterile saline and returned to each animal. After collecting the blood sample at t = 120 min, the mice were anesthetized with isoflurane and euthanized by cervical dislocation, and blood samples were collected from the heart to measure CRR hormones (30).

### Glucagon and NE injection

C57BL/6N male mice were bilaterally injected in the VMH with shPLA2 or shSCRM as described above. Three weeks later, these mice experienced RH for five days. On day 6, mice received i.p. injection of glucagon (0.1U/kg) in *ad libitum* fed condition. Blood glucose levels were measured at 0, 15, 30, 60, and 120 min after injection. On another day, the same mice received norepinephrine (1mg/kg) i.p. injection, and blood glucose levels were measured at the same time points following the injection.

### Measurement of serum hormones

After performing the clamp, the blood samples were centrifuged for 10 min at 1000 × *g* and maintained at −30°C until hormones were measured. The glucagon, corticosterone, epinepheline, and NE concentrations were measured with each ELISA kit (Enzo Life Sciences Inc., Farmingdale, NY). All the protocols followed the instructions provided by the kit.

### Immunohistochemistry (IHC)

C57BL/6 male mice received saline or ibuprofen (30 mg/kg body weight) *per os* (p.o.) using a gastric probe 30 min before i.p. injection of saline or insulin (0.75 U/kg). The mice were sacrificed using CO_2_ asphyxiation and perfused with heparinized saline transcardially at 60 min after i.p. injection. Brain sections (50 µm each) containing both VMH and ARC were collected. The floating sections were incubated with rabbit-anti-cFos antibody (1:1000, Cell Signaling Technology, #2250S, MA) in blocking solution (0.1 M phosphate buffer (PB) containing 4% normal guinea pig serum, 0.1% glycine, and 0.2% Triton X-100) overnight at room temperature. After rinsing with PB, the sections were incubated in a secondary antibody (1:500, Alexa 488 secondary antibody, Invitrogen, A11008, MA) for 2 h at room temperature. The stained sections were washed with PB 3 times and mounted on glass slides with DAPI Fluoromount-G (Southern Biotech, Birmingham, AL). Cells were automatically counted using an ImageJ software plugin (Analyze Particles).

### Statistical analysis

For repeated-measures analysis, two-way ANOVA was used when values over different times were analyzed, followed by the Sidak multiple comparisons tests. For the statistical analysis between multiple independent groups, one-way ANOVA followed by Tukey’s multiple comparisons tests. When only two groups were analyzed, statistical significance was determined by the unpaired Student’s *t*-test (two-tailed *p*-value). Prism 10 software (GraphPad Software, San Diego, CA) was used for these statistical analyses. A value of *p* < 0.05 was considered statistically significant. All data are shown as mean ± SEM.

## Results

### Glucose deprivation, induced by insulin and 2DG, decreases arachidonic acid-containing phospholipids in the hypothalamus in mice

To check the role of hypoglycemia in hypothalamic phospholipid metabolism, the signal intensity of 11 different phospholipids located in both the VMH and the ARC were measured at 30 min after the intraperitoneal (i.p.) insulin injection (Fig. 1a). After the insulin injection, blood glucose levels became 71.6 ± 6.5 mg/dL (Fig. 1b). Insulin injection decreased phosphatidylinositol (PI) (16:0/20:4), PI (18:0/20:4), PI (18:1/20:4), phosphatidylethanolamine (PE) (18:0/16:1), PE (18:0/20:4) and phosphatidylserine (PS) (18:0/16:0) compared to the saline group in the VMH (Fig. 1c-f) and PI (16:0/20:4), PI (18:0/20:4), and PI (18:1/20:4) in the ARC (Fig. 1c, g-i). We measured the activities of PLA1, PLA2, PLC, and PLD, which are enzymes that metabolize phospholipids. Insulin injection activated cPLA2 but not PLA1, secretory PLA2, PLC, and PLD in the hypothalamus (Fig. 1j-n).

**Fig. 1.**
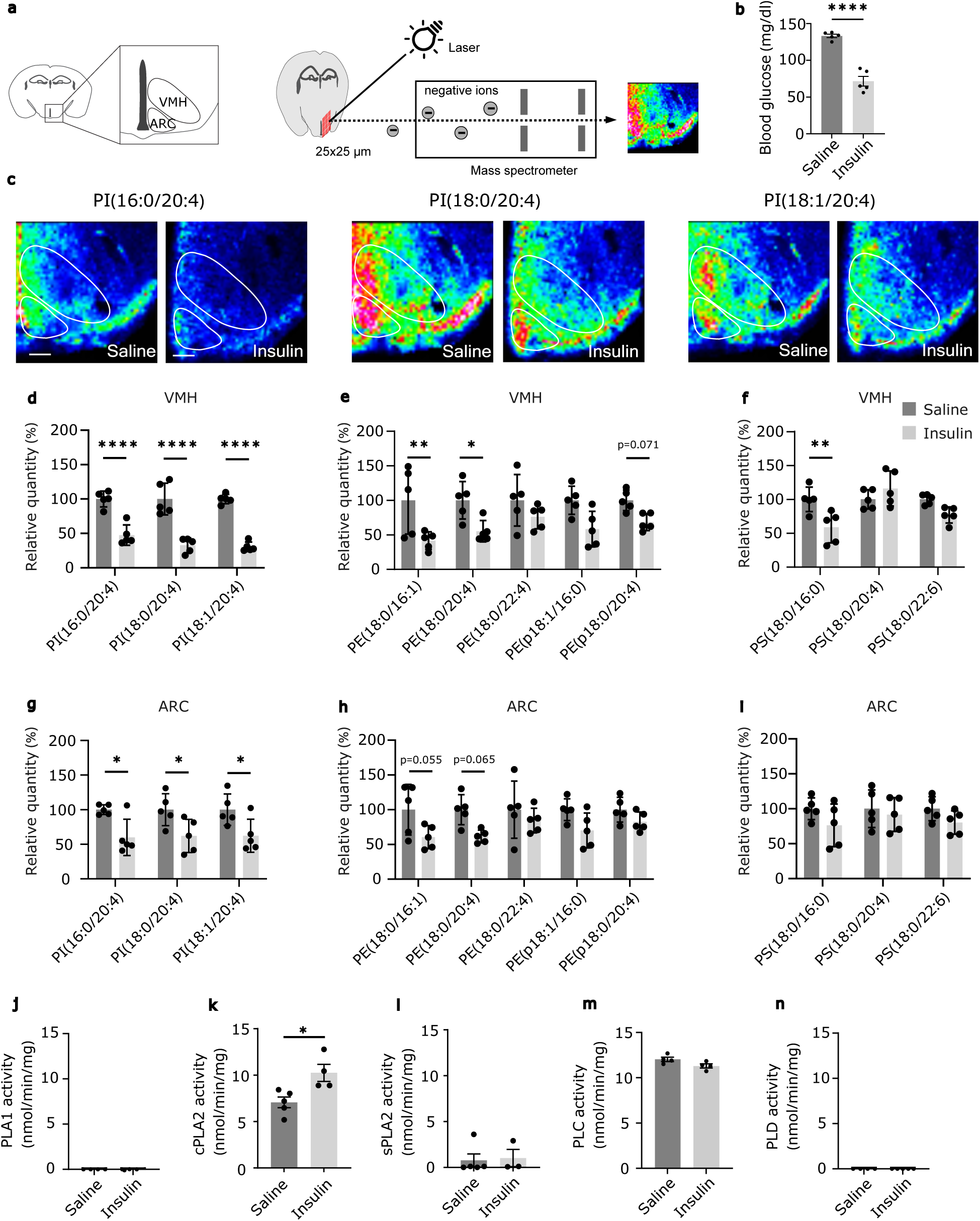
Insulin injection decreases phospholipids in the VMH. **a,** A schematic showing the ventromedial (VMH) and arcuate (ARC) nucleus of the hypothalamus, and imaging mass spectrometry (IMS). **b**, Blood glucose levels 30 min after the saline or insulin i.p. injection. **c**, Representative results of IMS showing the distribution of hypothalamic phosphatidylinositol (PI) (16:0/20:4), PI (18:0/20:4), and PI (18:1/20:4), in mice after injection with saline (left) or insulin (right). The white line shows the position of the VMH and ARC. Scale bar: 200 µm. **d-i,** Relative intensities of PI (d, g), phosphatidylethanolamine (PE) (e, h), and phosphatidylserine (PS) (f, i) in the VMH and ARC 30 min after the saline (n = 5) or insulin (n = 5) i.p. injection. **j-n,** Enzymatic activity of PLA1 (j), cytosolic PLA2 (k) and secretory PLA2 (l), PLC (m), and PLD (n) in the hypothalamus 30 min after the saline (n = 5) or insulin (n = 4) i.p. injection. All results were shown as mean ± SEM, \**p* < 0.05, \*\**p* < 0.01, \*\*\*\**p* < 0.0001. Student’s *t*-test (two-tailed *p*-value) and one-way ANOVA followed by Tukey’s multiple comparison tests were used for statistical analysis.

To further study the role of insulin in hypothalamic phospholipid metabolism, we used streptozotocin (STZ) to disrupt the pancreatic beta cells and insulin secretion. Blood glucose level became 442.8 ± 35.5 mg/dL 4 days after STZ treatment (Fig. 2a). When phospholipid intensity was examined in the VMH and ARC, no significant differences were observed in any types of phospholipids, including PI, PE, and PS in either hypothalamic nucleus (Fig. 2b-g). To evaluate the effect of hypoglycemia, we utilized 2-Deoxy-D-glucose (2DG), which cannot be processed in glycolysis, leading to glucose deprivation within the cell, which then activates CRRs. I.p. injection of 2DG increased blood glucose levels after 60 min (Fig. 2h). 2DG decreased the signal intensity of PI (18:0/20:4) and PE (p18:0/20:4) in the VMH compared to mice receiving a saline injection (Fig. 2i-k). 2DG injection also decreased PI (18:1/20:4), PE (p18:0/20:4) and PS (18:0/16:0) in the ARC (Fig. 2l-n). These data suggest that not only insulin but also hypoglycemia enhances arachidonic acid (AA)-containing phospholipid metabolism in the hypothalamus, at least in part.

**Fig. 2.**
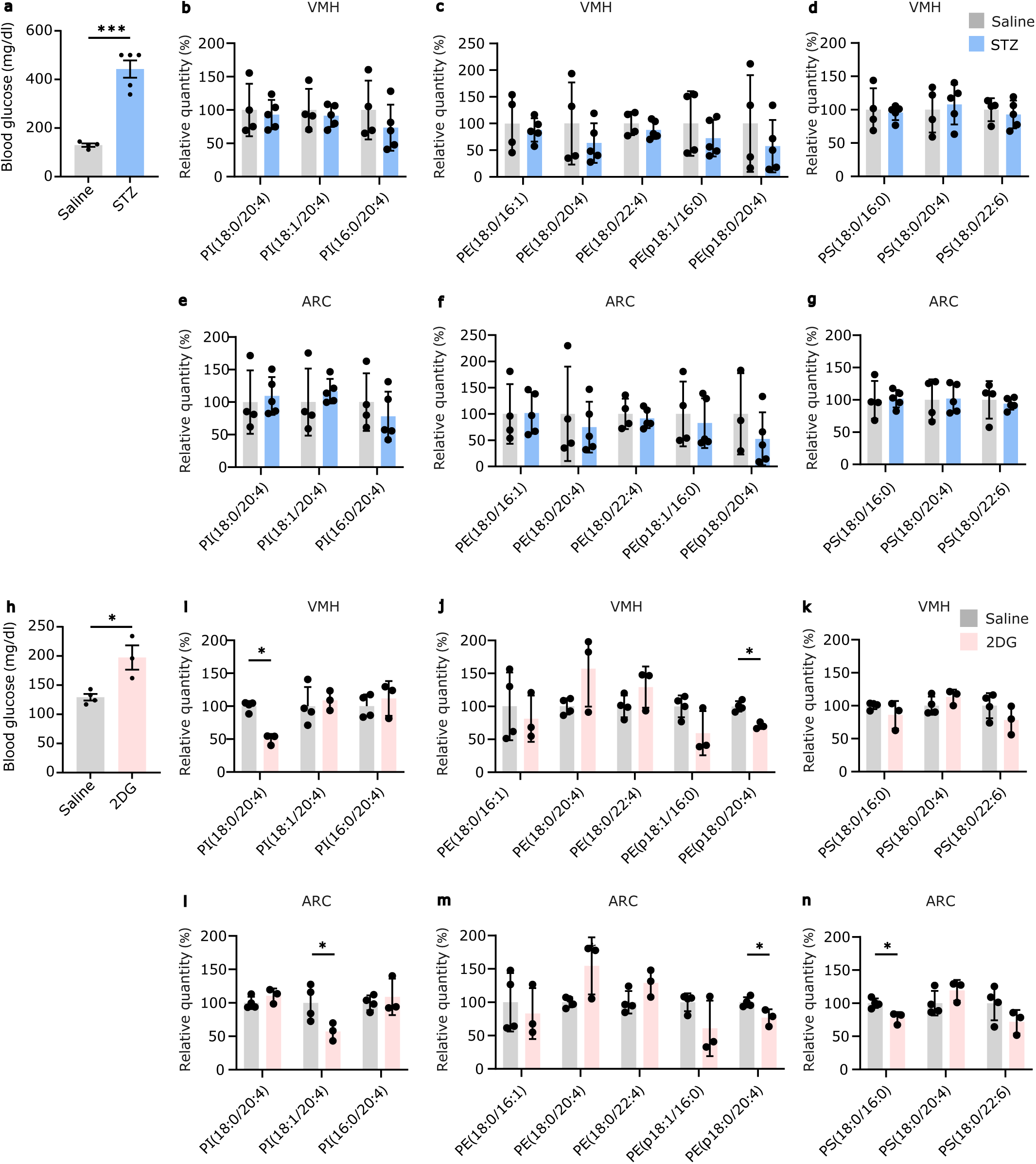
Glucose deprivation, but not insulin deficiency, changes phospholipids in the hypothalamus. **a**, Blood glucose levels 4 days after the saline or STZ injection. **b-g**, Relative intensities of phospholipids in the VMH (b-d) and ARC (e-g) 4 days after i.p. injection with saline (n = 4) or STZ (n = 5). **h**, Blood glucose levels in 30 min after the saline or 2DG i.p. injection. **i-n,** Relative intensities of phospholipids in the VMH (i-k) and ARC (l-n) 60 min after i.p. injection with saline (n = 4) or 2DG (n = 3). All results were shown as mean ± SEM, \**p* < 0.05. Student’s *t*-test (two-tailed *p*-value) and one-way ANOVA followed by Tukey’s multiple comparison tests were used for statistical analysis.

### Insulin increases prostaglandin production in the hypothalamus

AA is known to be used as a substrate of prostaglandins (PGs). Thus, we measured all types of PGs in the whole hypothalamus by LC-MS 30 min after insulin injection when blood glucose became 46.7 ± 9.8 mg/dL (Fig. 3a). Insulin injection increased the amount of 6-keto-PGF1α, 11-β-13,14-dihydro-15-keto-PGF2α, PGF2α, PGE2, and 20-hydroxy-PGF2α in the hypothalamus (Fig. 3b-g). Therefore, these results suggest that insulin injection induces AA release from phospholipids to produce PGs in the hypothalamus.

**Fig. 3.**
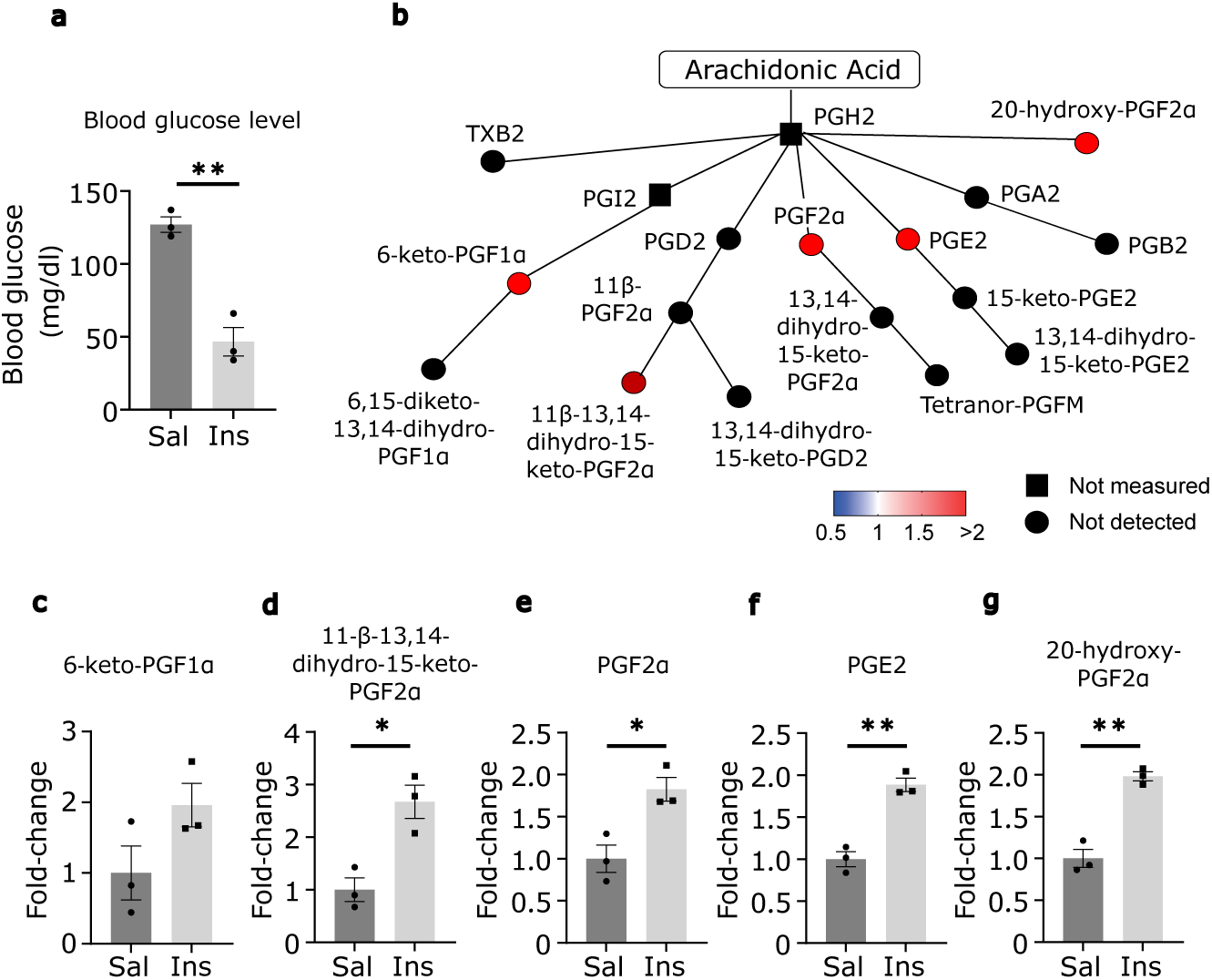
Insulin increases prostaglandins in the hypothalamus. **a,** Blood glucose levels 30 min after the saline or insulin i.p. injection. **b**, Relative amounts of hypothalamic prostaglandins mediated by cyclooxygenase showing mean fold-change in color. **c-g**, Fold-change of 6-keto-PGF1α (c), 11-β-13,14-dihydro-15-keto-PGF2α (d), PGF2α (e), PGF2 (f), and 20-hydroxy-PGF2α (g). All results were shown as mean ± SEM, \**p* < 0.05; \*\**p* < 0.01. The unpaired Student’s *t*-test (two-tailed *p*-value) was used for statistical analysis.

### Ibuprofen blocks CRRs and activity of GI neuron

To investigate the role of PGs in the brain in the regulation of systemic glucose metabolism, we injected ibuprofen i.c.v. and measured glucose production induced by i.p. injection of 2DG. 2DG injection activates CRRs and increased blood glucose levels (Fig. 4a, b). I.c.v. injection of ibuprofen lowered the 2DG-induced hyperglycemia (Fig. 4a, b). 2DG is also known to increase food intake. However, ibuprofen injection did not change feeding behavior compared to saline-injected mice (Fig. 4c).

**Fig. 4.**
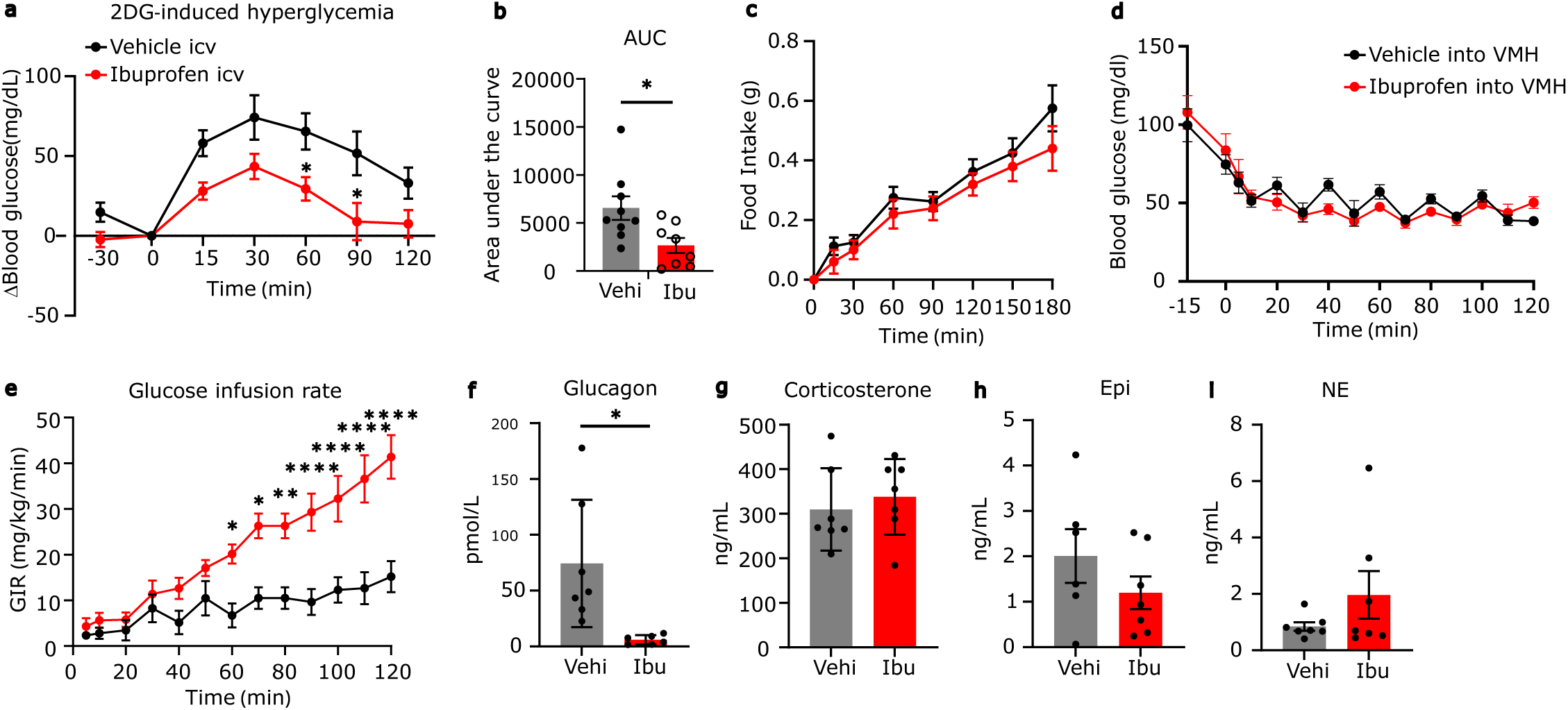
Central ibuprofen injection impairs glucose production in 2DG-induced glucose deprivation and hypoglycemic clamp. **a,** 2DG-induced hyperglycemia in mice that received ibuprofen or vehicle injection (i.c.v.). **b**, Area under the curve of blood glucose in (a). **c**, Food intake after i.p. injection of 2DG in ibuprofen or vehicle-injected mice (i.c.v.). **d,** Blood glucose levels during hyperinsulinemic-hypoglycemic clamp in mice that received ibuprofen or vehicle injection into the VMH bilaterally. Insulin infusion was started at t = 0 min. **e**, Glucose infusion rate to maintain hypoglycemia at 50 mg/dL during clamp. **f-i**, Concentration of serum hormones, glucagon (f), corticosterone (g), epinephrine (h), and norepinephrine (i), corrected at 120 min of the clamp experiment. All results were shown as mean ± SEM, \**p* < 0.05; \*\**p* < 0.01; \*\*\**p* < 0.001; \*\*\*\**p* < 0.0001. The unpaired Student’s *t*-test (two-tailed *p*-value) and two-way ANOVA followed by Sidak multiple comparison tests were used for statistical analysis.

To verify the role of PGs in the CRRs, we performed hyperinsulinemic-hypoglycemic clamp experiments in mice. In the hypoglycemic clamp, a constant rate of insulin (1 mU/kg/min) was infused into the circulation, while glucose was infused as needed to maintain blood glucose levels at 50 mg/dL (Fig. 4d). Ibuprofen injection into the VMH increased glucose infusion rate (GIR) (Fig. 4e), suggesting that ibuprofen inhibits glucose production during hypoglycemia and thus more glucose was needed to maintain at 50 mg/dL. Then, we measured CRRs-related hormones in the blood at 120 min of the clamp experiment. Ibuprofen injection significantly decreased glucagon levels but not corticosterone, epinephrine, and norepinephrine (NE) (Fig. 4f-i). These data suggest that hypothalamic PGs are necessary for glucagon secretion in response to hypoglycemia to induce glucose production. To measure the neuronal activity in the hypothalamic GI neurons that regulate CRRs, we measured cFos immunohistochemistry. Insulin (i.p.) significantly increased the number of cFos-positive cells in the ventrolateral part of the VMH (vlVMH), while oral injection of ibuprofen suppressed it (Fig. 5a, b). The number of cFos-positive cells was not altered by ibuprofen injection in the dorsomedial (dm) and central (c) part of the VMH and ARC (Fig. 5a-c).

**Fig. 5.**
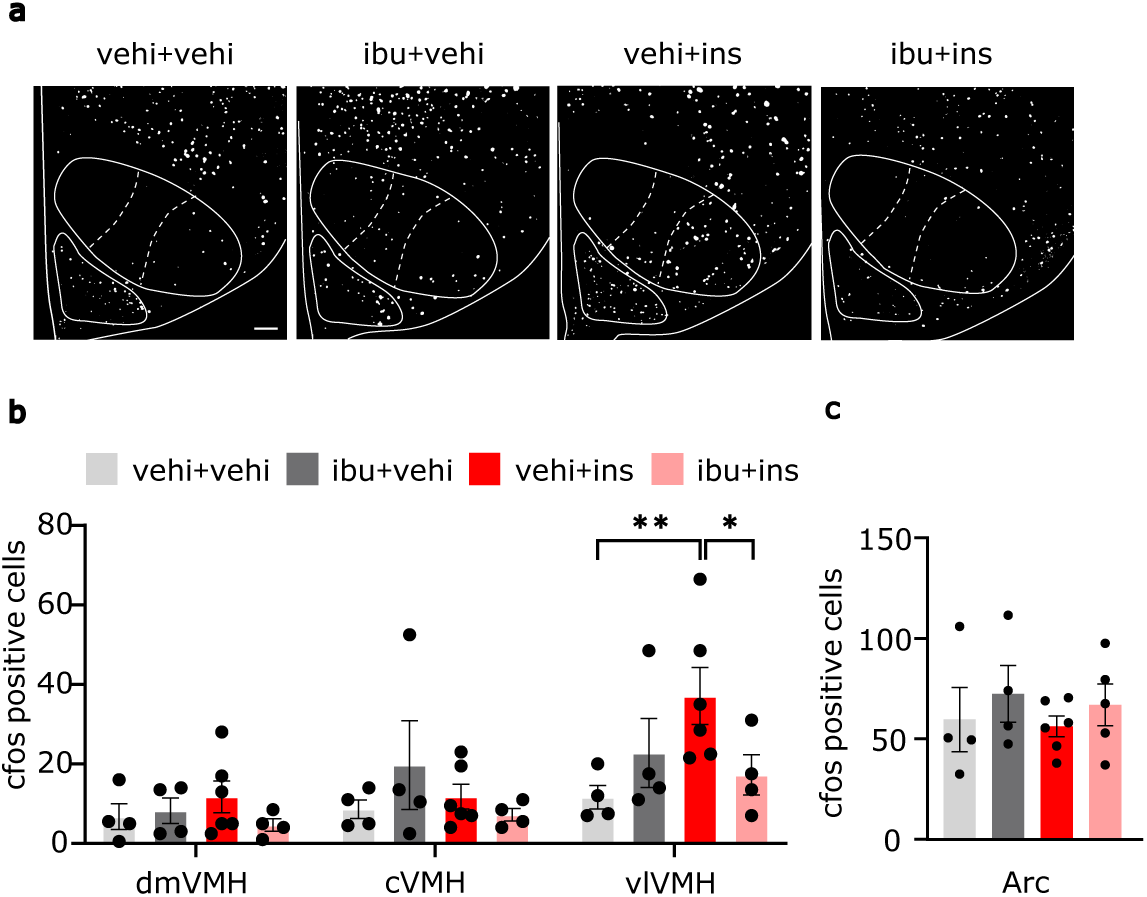
Ibuprofen decreases insulin-induced cFos-positive cells specifically in ventrolateral VMH. **a,** Representative micrographs of cFos immunoreactivity in the VMH and ARC. Scale bar: 100 µm. White lines indicate the boundary of the nucleus in the hypothalamus. **b-c,** Quantification of cFos cell number in dorsomedial (dmVMH), central (cVMH) and ventrolateral (vlVMH) part of the VMH (b) and arcuate nucleus (ARC) (c) in the mice that received vehicle (p.o.) and vehicle (i.p.) (vehi+vehi), ibuprofen (p.o.) and vehicle (i.p.) (ibu+vehi), vehicle (p.o.) and insulin (i.p.) (vehi+ins), ibuprofen (p.o.) and insulin (i.p.) (ibu+ins). All results were shown as mean ± SEM, \**p* < 0.05; \*\**p* < 0.01. One-way ANOVA followed by Tukey’s multiple comparison tests were used for statistical analysis.

### Hypothalamic PGs deteriorate glucose production after RH

To generate the RH mouse model, 2.5 U/kg of insulin was injected i.p. to induce hypoglycemia (40 mg/dL) once a day for 5 days (Fig. 6a, d, g). To inhibit the production of PGs in the hypothalamus, we injected AAV that expresses shRNA against cPLA2 (shPLA2) into the VMH (Fig. 6a), since cPLA2 is activated by insulin injection (Fig. 1k). AAV-infected cells were found in the vlVMH and ARC (Fig. 6b). cPLA2 mRNA in the hypothalamus was significantly decreased (Fig. 6c). On day 6, the hypoglycemic clamp was performed (Fig. 6a, e). In the control group, GIR was significantly increased in RH mice (Fig. 6f), suggesting that RH attenuated glucose production. shPLA2 group also received RH treatment for 5 days, and the hypoglycemic clamp was performed (Fig. 6g-i). GIR in the shPLA2 without RH was similar to that of control-RH mice and ibuprofen-injected mice (Fig. 4e, 6f), confirming that hypothalamic PG production enhances CRRs in the non-RH condition. However, the shPLA2 inhibited the increase in GIR after RH (Fig. 6i). RH decreased plasma glucagon and epinephrine at 120 min in the hypoglycemic clamp (Fig. 6j, k). RH did not change plasma norepinephrine, corticosterone, and growth hormone (Fig. 6l-n). shPLA2 did not improve glucagon, epinephrine, NE, corticosterone, or growth hormone levels (Fig. 6j-n). After RH, shPLA2 group had a higher glucose production after glucagon injection than shSCRM group (Fig. 6o), while NE-induced glucose production was comparable between groups (Fig. 6p).

**Fig. 6.**
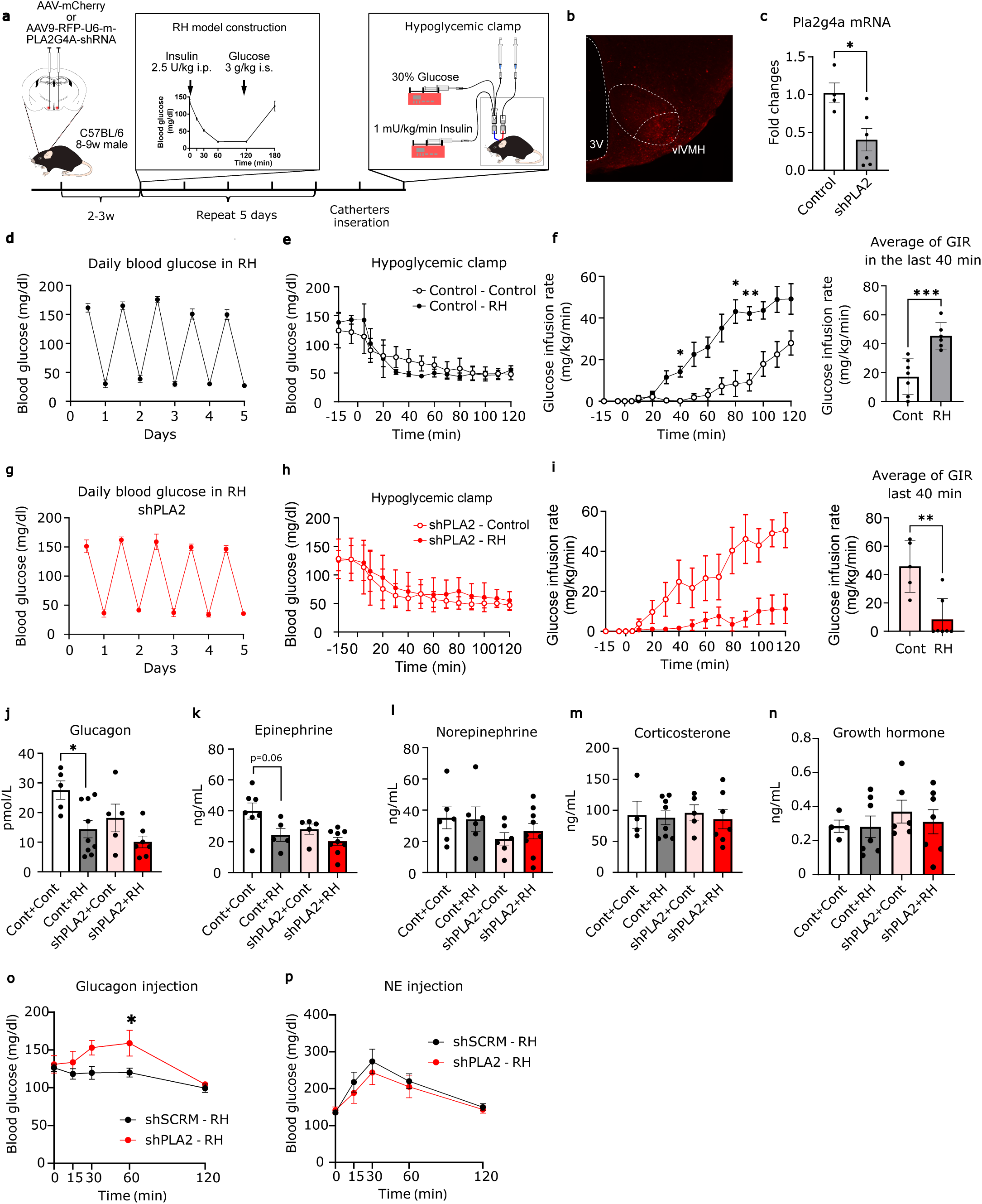
shRNA against cPLA2 in the hypothalamus improves RH-induced impairment of glucose production. **a**, A schematic illustrating AAV injection into the VMH and experimental timeline of AAV injection, RH, and hypoglycemic clamp. **b**, Representative image of RFP expression after AAV injection. **c**, mRNA expression of cPLA2 in the hypothalamus. **d,g**, Daily blood glucose levels during 5 days of recurrent hypoglycemia. **e,h**, Blood glucose levels during hypoglycemic clamp. Insulin infusion was started at t = 0 min. **f,i**, Glucose infusion rate to maintain hypoglycemia at 50 mg/dL during clamp and average GIR in the last 40 min. **j-n**, Concentration of serum hormones, glucagon (j), epinephrine (k), norepinephrine (l), corticosterone (m), and growth hormone (n), corrected at 120 min of the clamp experiment. **o-p**, Glucagon (o) and norepinephrine (p) sensitivities in shPLA2 or shSCRM injected mice after the RH. All results were shown as mean ± SEM, \**p* < 0.05; \*\**p* < 0.01; \*\*\**p* < 0.001; \*\*\*\**p* < 0.0001. The unpaired Student’s *t*-test (two-tailed *p*-value), one-way ANOVA followed by Tukey’s multiple comparison tests, and two-way ANOVA followed by Sidak multiple comparison tests were used for statistical analysis.

## Discussion

In this study, we found that 1) insulin, and partially insulin-induced hypoglycemia, increase PG production using arachidonic acid from phospholipids; 2) The PG production in the VMH is necessary for the neuronal activation in the vlVMH, glucagon secretion from the pancreas, and an increase in hepatic glucose production during acute hypoglycemia; 3) RH attenuates glucagon secretion and glucose production; 4) cPLA2 in the hypothalamus is critical for attenuating glucose production after RH by changing glucagon sensitivity.

NSAIDs, including salicylates, were used to treat diabetic patients in the early 1900s (31). NSAIDs cause gastrointestinal bleeding (31) and severe hypoglycemia when used with other blood glucose-lowering agents (32). Hence, they are no longer used as anti-diabetic drugs. The mechanism of the hypoglycemic effect of NSAIDs has been studied only in the peripheral tissues (26,27). While NSAIDs are the most commonly used drug for analgesics due to their inhibition of PG production in the brain, research on their effect on lowering blood glucose levels in the central nervous system is limited. I.c.v. injection of PGD2, PGE1, PGE2, or PGF2α increases the glucose concentration in the hepatic vein (33). Central PGF2α has the most substantial effect but is mediated by epinephrine secretion from the adrenal medulla, not glucagon (33).

The VMH plays a vital role in the regulation of CRRs to hypoglycemia. A lesion of the VMH suppresses CRRs in response to systemic injection of 2DG (34). 2DG perfusion in the VMH via microdialysis increases glucagon, epinephrine, and NE secretion (35). In contrast, glucose perfusion in the VMH during the hypoglycemic clamp blocks increases in the secretion of CRR-related hormones (36). Synaptic glutamate release from the VMH-specific steroidogenic factor 1 (SF1) neuron regulates CRRs (37). Neuronal pathways from the PBN to the VMH (10) and from VMH to the BNST regulate CRRs (38). Therefore, it is considered that the VMH has a critical role in detecting low blood glucose levels and changing CRRs. In our previous paper (17), the knockdown of cPLA2 in SF1 neurons did not change blood glucose levels after systemic 2DG injection compared to control mice. Therefore, neurons in the dmVMH, including SF1 neurons, may not be necessary for NSAID-related change in CRRs. Conversely, estrogen receptor alpha (ERα)-expressing neurons in the vlVMH are involved in the CRRs (39). Both GI-type ERα neurons projecting to the medioposterior ARC and GE-type ERα neurons projecting to the dorsal raphe nucleus regulate CRRs (39). In the present data, we found that ibuprofen affects neuronal activity in the vlVMH but not dmVMH and cVMH. Thus, neuronal circuits from ERα neurons in the vlVMH projecting to the medioposterior ARC may regulate NSAID-related hypoglycemia.

RH is a critical incident in the treatment of diabetes, and it can result in hypoglycemia-associated autonomic failure. RH increases hexokinase activity and attenuates the activity of glucose-sensing neurons in the VMH (40). Indeed, we found that the RH decreased plasma glucagon, epinephrine, and glucose production during the hypoglycemic clamp. Expression of shPLA2 in the VMH improved the RH-induced attenuation of glucose production. However, shPLA2 did not change glucagon or epinephrine production. Instead, it enhanced glucagon sensitivity to increase glucose levels. Liu Z et al. reported that RH changes adrenergic sensitivity in the liver and visceral fat (20). In their RH model rat, RH decreased epinephrine secretion, but it did not attenuate glucagon secretion. Differences among species may exist in RH-induced changes; however, rats and mice show varying sensitivities to adrenaline and glucagon, which had their secretions reduced in each rodent, respectively. Few papers studied on glucagon sensitivity. Glucagon sensitivity is not altered in type 1 diabetes mellitus (T1DM) (41). Metabolic-associated steatotic liver disease (MASLD) and MASLD-related gestational diabetes mellitus impair glucagon sensitivity (42,43). Conversely, pancreatectomised patients increase hepatic glucagon sensitivity (44). In mice, western diet-fed mice, high-fat diet-fed, and db/db mice do not change glucagon sensitivity (45). However, there are no reports of the brain modulating glucagon sensitivity as far as we searched.

Recently, the importance of glucagon in both T1 and T2DM is apparent (46). In our study, intra-hypothalamic injection of ibuprofen and experiencing RH decreased glucagon secretion during hypoglycemia. In T2DM patients, NSAIDs decrease blood glucose concentration and HbA1c (47,48). In a retrospective cohort study, NSAIDs also reduced the risk of becoming T2DM (49). Since glucagon concentrations were not measured in these papers, the relationship between NSAIDs, glucagon, and T2DM is still unclear. Further study is needed to clarify the mechanisms by which NSAIDs improve T1 and T2DM. The possibility that the brain regulates glucagon sensitivity may be an interesting research topic and has the potential to open a new field of glucagon research.

The present study revealed the importance of PGs in the hypothalamus for the recovery from hypoglycemia. Hypothalamic PGs regulate the activity of GI neurons and enhance glucagon secretion. RH impairs glucagon and epinephrine secretion. Hypothalamic PGs attenuate glucose production after RH by changing glucagon sensitivity. More studies are needed to broaden our understanding of strategies to prevent RH.

## Article Information

## Acknowledgments

We thank Nur Farehan Asgar, PhD, for editing a draft of this manuscript. This work was supported by JSPS KAKENHI (Grant Number 21H02352, 21K18275T, 23H00512); Japan Agency for Medical Research and Development (AMED-RPIME, Grant Number JP21gm6510009h0001, JP22gm6510009h9901, 23gm6510009h9903, 24gm6510009h9904); the Takeda Science Foundation; the Uehara Memorial Foundation; Astellas Foundation for Research on Metabolic Disorders; Suzuken Memorial Foundation; Akiyama Life Science Foundation; Narishige Neuroscience Research Foundation; JST SPRING (JPMJSP2119); and Daiichi Sankyo Foundation of Life Science.

## Author contributions

C.T. conceived this study, designed the experiments, and supervised the entire study. T.A. and S. X. performed most of the experiments and analysis. Y.S. performed LC-MS measurements. C.T. and T.H. performed imaging mass spectrometry. M.L., T.I. and Y.A. performed 2DGTT and hypoglycemic clamp. N.W. and Z.N. performed IHC. T.A., S.X., Y.A., C.T. wrote the manuscript. N.I. and S.D. assisted in preparing the manuscript.

---

**Table S1.**
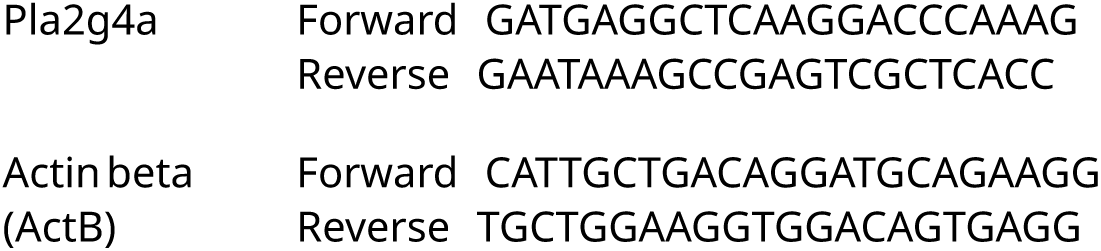
Mouse qPCR Primer sequences.

